# Exploring the single-cell RNA-seq analysis landscape with the scRNA-tools database

**DOI:** 10.1101/206573

**Authors:** Luke Zappia, Belinda Phipson, Alicia Oshlack

**Affiliations:** Bioinformatics, Murdoch Children’s Research Institute, Melbourne, Victoria, Australia; School of Biosciences, Faculty of Science, University of Melbourne, Melbourne, Victoria, Australia

## Abstract

As single-cell RNA-sequencing (scRNA-seq) datasets have become more widespread the number of tools designed to analyse these data has dramatically increased. Navigating the vast sea of tools now available is becoming increasingly challenging for researchers. In order to better facilitate selection of appropriate analysis tools we have created the scRNA-tools database (www.scRNA-tools.org) to catalogue and curate analysis tools as they become available. Our database collects a range of information on each scRNA-seq analysis tool and categorises them according to the analysis tasks they perform. Exploration of this database gives insights into the areas of rapid development of analysis methods for scRNA-seq data. We see that many tools perform tasks specific to scRNA-seq analysis, particularly clustering and ordering of cells. We also find that the scRNA-seq community embraces an open-source approach, with most tools available under open-source licenses and preprints being extensively used as a means to describe methods. The scRNA-tools database provides a valuable resource for researchers embarking on scRNA-seq analysis and records of the growth of the field over time.

**Author summary:** In recent years single-cell RNA-sequeing technologies have emerged that allow scientists to measure the activity of genes in thousands of individual cells simultaneously. This means we can start to look at what each cell in a sample is doing instead of considering an average across all cells in a sample, as was the case with older technologies. However, while access to this kind of data presents a wealth of opportunities it comes with a new set of challenges. Researchers across the world have developed new methods and software tools to make the most of these datasets but the field is moving at such a rapid pace it is difficult to keep up with what is currently available. To make this easier we have developed the scRNA-tools database and website (www.scRNA-tools.org). Our database catalogues analysis tools, recording the tasks they can be used for, where they can be downloaded from and the publications that describe how they work. By looking at this database we can see that developers have focued on methods specific to single-cell data and that they embrace an open-source approach with permissive licensing, sharing of code and preprint publications.

## Introduction

Single-cell RNA-sequencing (scRNA-seq) has rapidly gained traction as an effective tool for interrogating the transcriptome at the resolution of individual cells. Since the first protocols were published in 2009 [1] the number of cells profiled in individual scRNA-seq experiments has increased exponentially, outstripping Moore’s Law [2]. This new kind of transcriptomic data brings a demand for new analysis methods. Not only is the scale of scRNA-seq datasets much greater than that of bulk experiments but there are also a variety of challenges unique to the single-cell context [3]. Specifically, scRNA-seq data is extremely sparse (there is no expression measured for many genes in most cells), it can have technical artefacts such as low-quality cells or differences between sequencing batches and the scientific questions of interest are often different to those asked of bulk RNA-seq datasets. For example many bulk RNA-seq datasets are generated to detect differentially expressed genes through a designed experiment while many scRNA-seq experiments aim to identify or classify cell types in complex tissues.

The bioinformatics community has embraced this new type of data at an astonishing rate, designing a plethora of methods for the analysis of scRNA-seq data. As such, keeping up with the current state of scRNA-seq analysis is now a significant challenge as the field is presented with a huge number of choices for analysing a dataset. Since September 2016 we have collated and categorised scRNA-seq analysis tools as they have become available. This database is being continually updated and is publicly available at www.scRNA-tools.org. In order to help researchers navigate the vast ocean of analysis tools we categorise tools in the database in the context of the typical phases of an scRNA-seq analysis. Through the analysis of this database we show trends in not only the analysis applications these methods address but how they are published, licensed and the platforms they use. Based on this database we gain insight into the current state of current tools in this rapidly developing field.

## Overview of the scRNA-tools database

The scRNA-tools database contains information on software tools specifically designed for the analysis of scRNA-seq data. For a tool to be eligible to be included in the database it must be available for download and public use. This can be from a software package repository (such as Bioconductor [4], CRAN or PyPI), a code sharing website such as GitHub or directly from a private website. Various details of the tools are recorded such as the programming language or platform they use, details of any related publications, links to the source code and the associated software license. Tools are also categorised according to the analysis tasks they are able to perform. Most tools are added after a preprint or publication becomes available but some have been added after being mentioned on social media or in similar collections such as Sean Davis’ awesome-single-cell page (https://github.com/seandavi/awesome-single-cell).

The scRNA-tools website provides a profile for each tool, with links to publications and code repositories, as well as an index by analysis category. Both of these pages can be sorted in a variety of ways, including by the number of associated publications or citations. We also provide an interactive table that allows users to filter and sort tools in more sophisticated ways to find those most relevant to their needs. An additional page shows live and up-to-date versions of some of the analysis and visualisations of the database presented below. We welcome contributions to the database from the wider community via submitting an issue to the project GitHub page (https://github.com/Oshlack/scRNA-tools) or by filling in the submission form on the scRNA-tools website.

When the database was first constructed it contained 70 scRNA-seq analysis tools representing the majority of work in the field during the three years from the publication of SAMstrt [5] in November 2013 up to September 2016. In the time since then over 130 new tools have been added (Fig 1A). The almost tripling of the number of available tools in such a short time demonstrates the booming interest in scRNA-seq and its maturation from a technique requiring custom-built equipment with specialised protocols to a commercially available product.

**Fig 1.**
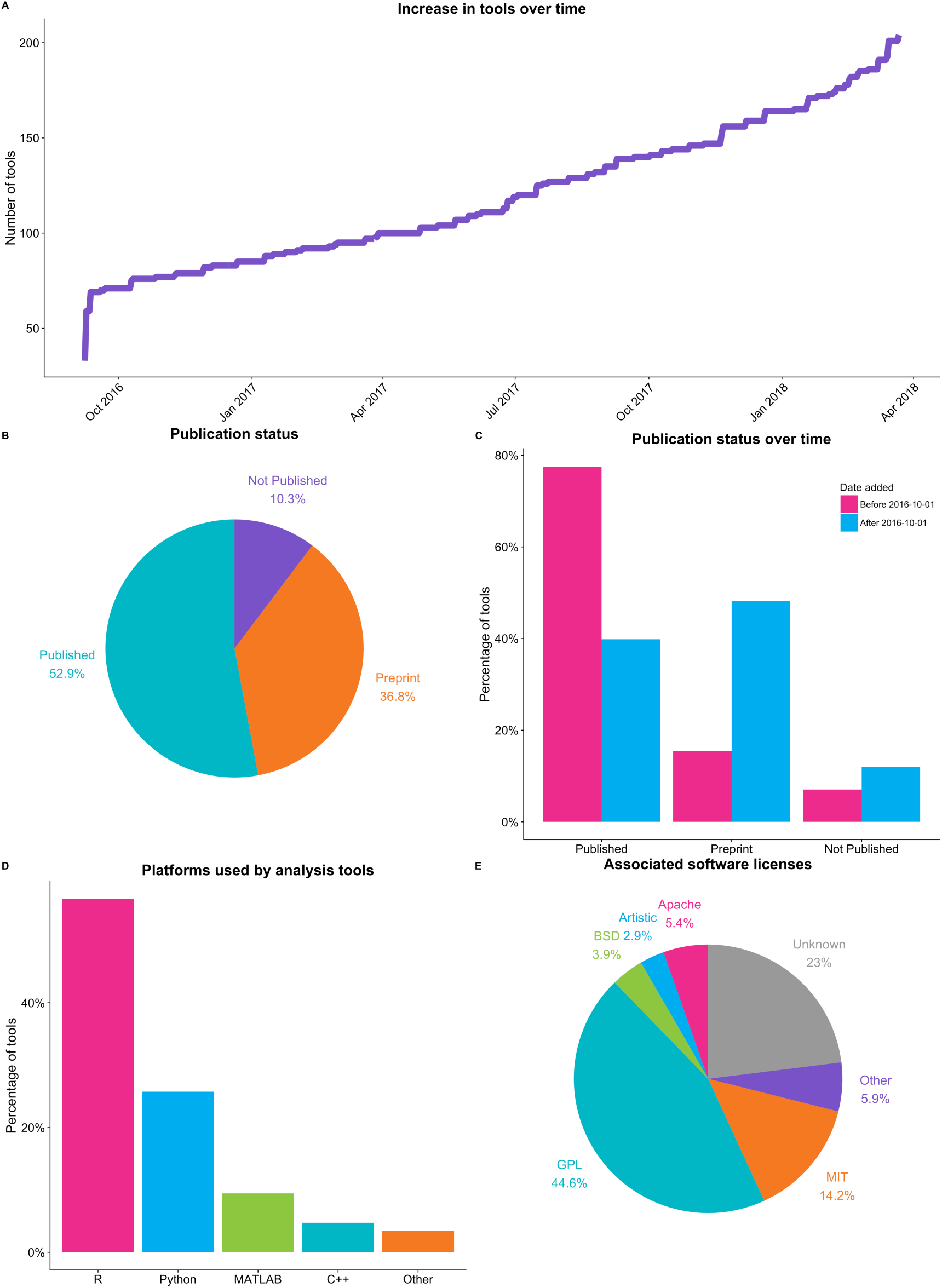
(A) Number of tools in the scRNA-tools database over time. Since the scRNA-seq tools database was started in September 2016 more than 130 new tools have been released. (B) Publication status of tools in the scRNA-tools database. Over half of the tools in the full database have at least one published peer-revirew paper while another third are described in preprints. (C) When stratified by the date tools were added to the database we see that the majority of tools added before October 2016 are published, while around half of newer tools are available only as preprints. Newer tools are also more likely to be unpublished in any form. (D) The majority of tools are available using either the R or Python programming languages. (E) Most tools are released under a standard open-source software license, with variants of the GNU Public License (GPL) being the most common. However licenses could not be found for a large proportion of tools. Up-to-date versions of these plots (with the exception of C) are available on the analysis page of the scRNA-tools website.

### Publication status

Most tools have been added to the scRNA-tools database after coming to our attention in a publication or preprint describing their method and use. Of all the tools in the database about half have at least one publications in a peer-reviewed journal and another third are described in preprint articles, typically on the bioRxiv preprint server (Fig 1B). Tools can be split into those that were available when the database was created and those that have been added since. We can see that the majority of older tools have been published while more recent tools are more likely to only be available as preprints (Fig 1C). This is a good demonstration of the delay imposed by the traditional publication process. By publishing preprints and releasing software via repositories such as GitHub scRNA-seq tool developers make their tools available to the community much earlier, allowing them to be used for analysis and their methods improved prior to formal publication [6].

### Platforms and licensing

Developers of scRNA-seq analysis tools have choices to make about what platforms they use to create their tools, how they make them available to the community and whether they share the source code. We find that the most commonly used platform for creating scRNA-seq analysis tools is the R statistical programming language, with many tools made available through the Bioconductor or CRAN repositories (Fig 1D). Python is the second most popular language, followed by MATLAB, a proprietary programming language, and the lower-level C++. The use of R and Python is consistent with their popularity across a range of data science fields. In particular the popularity of R reflects its history as the language of choice for the analysis of bulk RNA-seq datasets and a range of other biological data types.

The majority of tools in the scRNA-tools database have chosen to take an open-source approach, making their code available under permissive licenses (Fig 1E). We feel this reflects the general underlying sentiment and willingness of the bioinformatics community to share and build upon the work of others. Variations of the GNU Public License (GPL) are the most common, covering almost half of tools. This license allows free use, modification and distribution of source code, but also has a “copyleft” nature which requires any derivatives to disclose their source code and use the same license. The MIT license is the second most popular which also allows use of code for any purpose but without any restrictions on distribution or licensing. The appropriate license could not be identified for almost a quarter of tools. This is problematic as fellow developers must assume that source code cannot be reused, potentially limiting the usefulness of the methods in those tools. Tool owners are strongly encouraged to clearly display their license in source code and documentation to provide certainty to the community as to any restrictions on the use of their work.

## Categories of scRNA-seq analysis

Single-cell RNA-sequencing is often used to explore complex mixtures of cell types in an unsupervised manner. As has been described in previous reviews a standard scRNA-seq analysis in this setting consists of several tasks which can be completed using various tools [7–11]. In the scRNA-tools database we categorise tools based on the analysis tasks they perform. Here we group these tasks into four broad phases of analysis: data acquisition, data cleaning, cell assignment and gene identification (Fig 2). The data acquisition phase (Phase 1) takes the raw nucleotide sequences from the sequencing experiment and returns a matrix describing the expression of each gene (rows) in each cell (columns). This phase consists of tasks common to bulk RNA-seq experiments, such as alignment to a reference genome or transcriptome and quantification of expression, but is often extended to handle Unique Molecular Identifiers (UMIs) [12]. Once an expression matrix has been obtained it is vital to make sure the resulting data is of high enough quality. In the data cleaning phase (Phase 2) quality control of cells is performed as well as filtering of uninformative genes. Additional tasks may be performed to normalise the data or impute missing values. Exploratory data analysis tasks are often performed in this phase, such as viewing the datasets in reduced dimensions to look for underlying structure.

**Fig 2.**
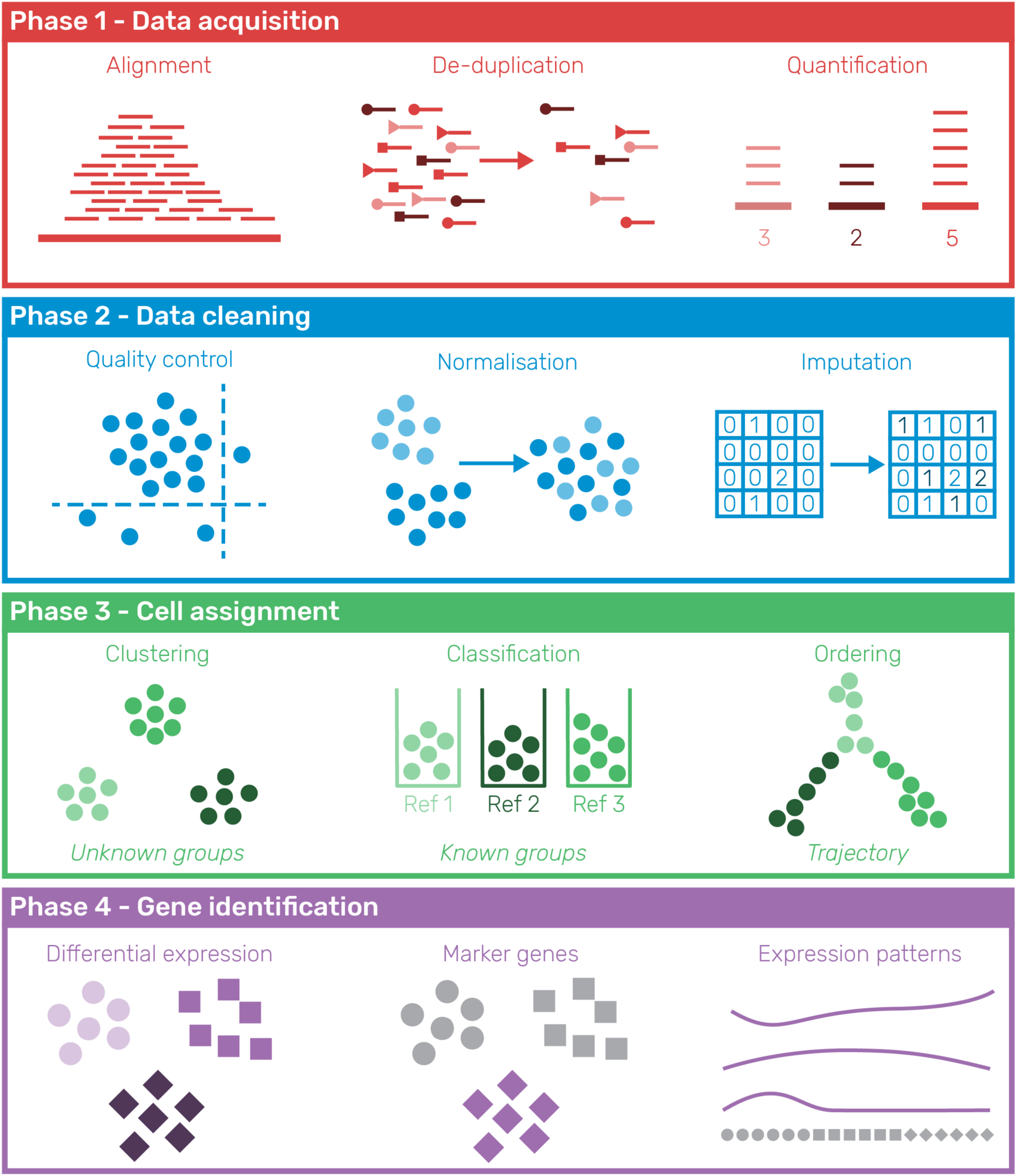
Phases of a typical unsupervised scRNA-seq analysis process. In Phase 1 (data acquisition) raw sequencing reads are converted into a gene by cell expression matrix. For many protocols this requires the alignment of genes to a reference genome and the assignment and de-duplication of Unique Molecular Identifiers (UMIs). The data is then cleaned (Phase 2) to remove low-quality cells and uninformative genes, resulting in a high-quality dataset for further analysis. The data can also be normalised and missing values imputed during this phase. Phase 3 assigns cells, either in a discrete manner to known (classification) or unknown (clustering) groups or to a position on a continuous trajectory. Interesting genes (eg. differentially expressed, markers, specific patterns of expression) are then identified to explain these groups or trajectories (Phase 4).

The high-quality expression matrix is the focus of the next phases of analysis. In Phase 3 cells are assigned, either to discrete groups via clustering or along a continuous trajectory from one cell type to another. As high-quality reference datasets become available it will also become feasible to classify cells directly into different cell types. Once cells have been assigned attention turns to interpreting what those assignments mean. Identifying interesting genes (Phase 4), such as those that are differentially expressed across groups, marker genes expressed in a single group or genes that change expression along a trajectory, is the typical way to do this. The biological significance of those genes can then be interpreted to give meaning to the experiment, either by investigating the genes themselves or by getting a higher level view through techniques such as gene set testing.

While there are other approaches that could be taken to analyse scRNA-seq data these phases represent the most common path from raw sequencing reads to biological insight applicable to many studies. An exception to this may be experiments designed to test a specific hypothesis where cell populations may have been sorted or the interest lies in differences between experimental conditions rather than cell types. In this case Phase 3 may not be required and slightly different tools or approaches may be used by many of the same challenges will apply. In addition, as the field expands and develops it is likely that data will be used in new ways to answer other biological questions, requiring new analysis techniques. Descriptions of the categories in the scRNA-tools database are given in Table 1, along with the associated analysis phases.

**Table 1.** Descriptions of categories for tools in the scRNA-tools database

| Phase | Category | Description |
| --- | --- | --- |
| Phase 1 | Alignment | Alignment of sequencing reads to a reference |
| Phase 1 | Assembly | Tools that perform assembly of scRNA-seq reads |
| Phase 1 | UMIs | Processing of Unique Molecular Identifiers |
| Phase 1 | Quantification | Quantification of expression from reads |
| Phase 2 | Quality Control | Removal of low-quality cells |
| Phase 2 | Gene Filtering | Removal of lowly expressed or otherwise<br>uninformative genes |
| Phase 2 | Imputation | Estimation of expression where zeros have been<br>observed |
| Phase 2 | Normalisation | Removal of unwanted variation that may affect<br>results |
| Phase 2 | Cell Cycle | Assignment or correction of stages of the cell cycle,<br>or other uses of cell cycle genes, or genes associated<br>with similar processes |
| Phase 3 | Classification | Assignment of cell types based on a reference dataset |
| Phase 3 | Clustering | Unsupervised grouping of cells based on expression profiles |
| Phase 3 | Ordering | Ordering of cells along a trajectory |
| Phase 3 | Rare Cells | Identification of rare cell populations |
| Phase 3 | Stem Cells | Identification of cells with stem-like characteristics |
| Phase 4 | Differential Expression | Testing of differential expression across groups of cells |
| Phase 4 | Expression Patterns | Detection of genes that change expression across a trajectory |
| Phase 4 | Gene Networks | Identification or use of co-regulated gene networks |
| Phase 4 | Gene Sets | Testing for over representation or other uses of annotated gene sets |
| Phase 4 | Marker Genes | Identification or use of genes that mark cell populations |
| Multiple | Dimensionality Reduction | Projection of cells into a lower dimensional space |
| Multiple | Interactive | Tools with an interactive component or a graphical user interface |
| Multiple | Variable Genes | Identification or use of highly (or lowly) variable genes |
| Multiple | Visualisation | Functions for visualising some aspect of scRNA-seq data or analysis |
| Other | Allele Specific | Detection of allele-specific expression |
| Other | Alternative Splicing | Detection of alternative splicing |
| Other | Haplotypes | Use or assignment of haplotypes |
| Other | Immune | Assignment of receptor sequences and immune cell clonality |
| Other | Integration | Combining of scRNA-seq datasets or integration with other single-cell data types |
| Other | Modality | Identification or use of modality in gene expression |
| Other | Simulation | Generation of synthetic scRNA-seq datasets |
| Other | Transformation | Transformation between expression levels and some other measure |
| Other | Variants | Detection or use of nucleotide variants |

### Trends in scRNA-seq analysis tasks

Each of the tools in the database is assigned to one or more analysis categories. We investigated these categories in further detail to give insight into the trends in scRNA-seq analysis. Fig 3A shows the frequency of tools performing each of the analysis tasks. Visualisation is the most commonly included task and is important across all stages of analysis for exploring and displaying data and results. Tasks for assigning cells (ordering and clustering) are the next most common. This has been the biggest area of development in single-cell analysis with clustering tools such as Seurat [[13]; Butler2017-lb], SC3 [14] and BackSPIN [15] being used to identify cell types in a sample and trajectory analysis tools (for example Monocle [[16]; Qiu2017-bi; Qiu2017-mq], Wishbone [17] and DPT [18]) being used to investigate how genes change across developmental processes. These areas reflect the new opportunities for analysis provided by single-cell data that are not possible with bulk RNA-seq experiments.

**Fig 3.**
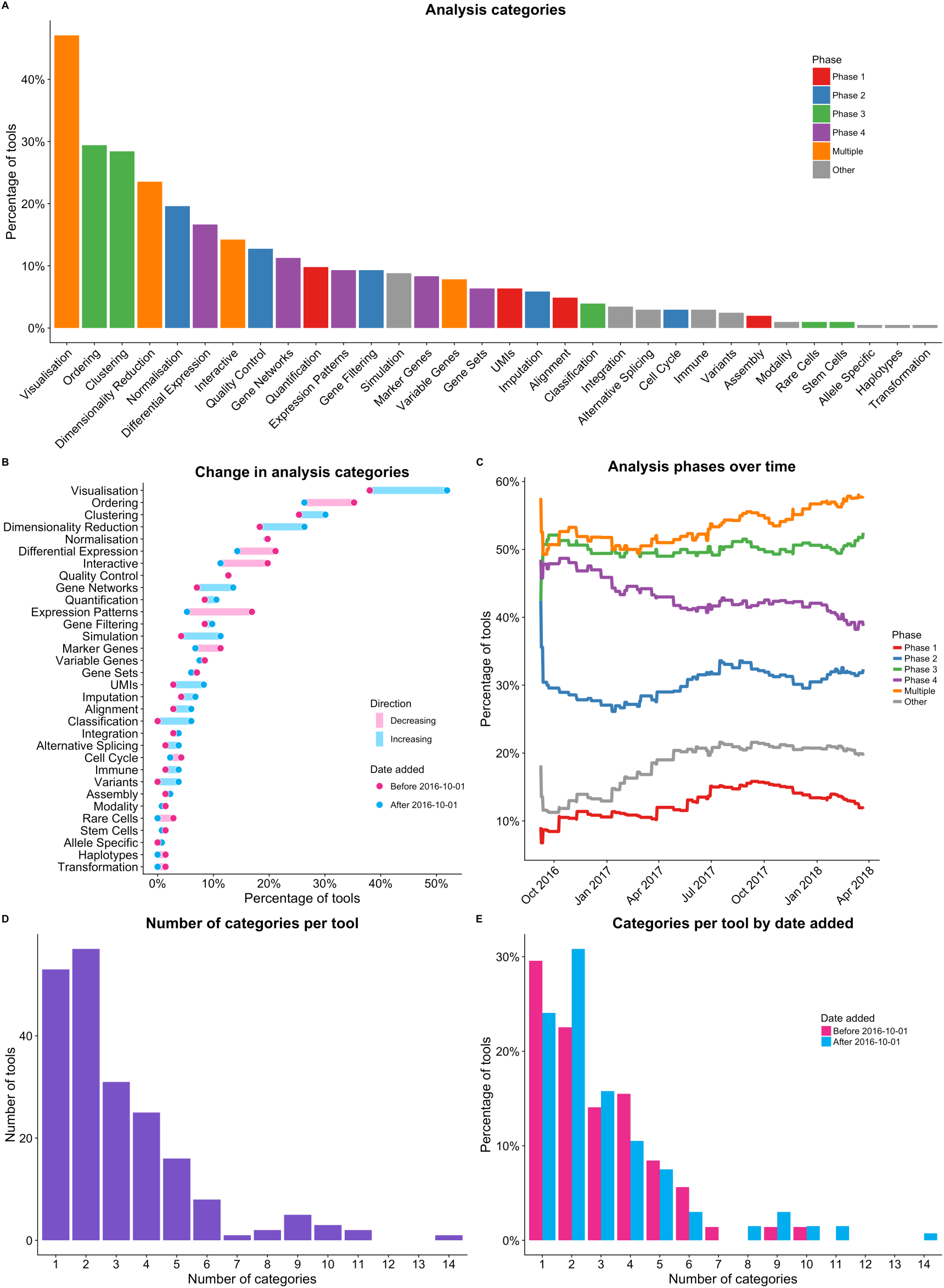
(A) Categories of tools in the scRNA-tools database. Each tool can be assigned to multiple categories based on the tasks it can complete. Categories associated with multiple analysis phases (visualisation, dimensionality reduction) are among the most common, as are categories associated with the cell assignment phase (ordering, clustering). (B) Changes in analysis categories over time, comparing tools added before and after October 2016. There have been significant increases in the percentage of tools associated with visualisation, dimensionality reduction, gene networks and simulation. Categories including expression patterns, ordering and interactivity have seen relative decreases. (C) Changes in the percentage of tools associated with analysis phases over time. The percentage of tools involved in the data acquisition and data cleaning phases have increased, as have tools designed for alternative analysis tasks. The gene identification phase has seen a relative decrease in the number of tools. (D) The number of categories associated with each tools in the scRNA-tools database. The majority of tools perform few tasks. (E) Most tools that complete many tasks are relatively recent.

Dimensionality reduction is also a common task and has applications in visualisation (via techniques such as t-SNE [19]), quality control and as a starting point for analysis. Testing for differential expression (DE) is perhaps the most common analysis performed on bulk RNA-seq datasets and it is also commonly applied by many scRNA-seq analysis tools, typically to identify genes that are different in one cluster of cells compared to the rest. However it should be noted that the DE testing applied by scRNA-seq tools is often not as sophisticated as the rigorous statistical frameworks of tools developed for bulk RNA-seq such as edgeR [20,21], DESeq2 [22] and limma [23], often using simple statistical tests such as the likelihood ratio test. While methods designed to test DE specifically in single-cell datasets do exist (such as SCDE [24], and scDD [25]) it is still unclear whether they improve on methods that have been established for bulk data [26–28], with the most comprehensive comparison to date finding that bulk methods do not perform significantly worse than those designed for scRNA-seq data [29].

To investigate how the focus of scRNA-seq tool development has changed over time we again divided the scRNA-tools database into tools added before and after October 2016. This allowed us to see which analysis tasks are more common in recently published tools. We looked at the percentage of tools in each time period that performed tasks in the different analysis categories (Fig 3B). Some categories show little change in the proportion of tools that perform them while other areas have changed significantly. Specifically, both visualisation and dimensionality reduction are more commonly addressed by recent tools. The UMIs category has also seen a big increase recently as UMI based protocols have become commonly used and tools designed to handle the extra processing steps required have been developed (e.g. UMI-tools [30], umis [31], zUMIs [32]). Simulation is a valuable technique for developing, testing and validating scRNA-seq tools. More packages are now including their simulation functions and some tools have been developed for the specific purpose of generating realistic synthetic scRNA-seq datasets (e.g. powsimR [33], Splatter [34]). Classification of cells into known groups has also increased as reference datasets become available and more tools are identifying or making use of co-regulated gene networks.

Some categories have seen a decrease in the proportion of tools they represent, most strikingly testing for changes expression patterns along a trajectory. This is likely related to the change in cell ordering analysis which is the focus of a lower percentage of tools added after October 2016. The ordering of cells along a trajectory was one of the first developments in scRNA-seq analysis and a decrease in the development of these tools could indicate that researchers have moved on to other techniques or that use has converged on a set of mature tools.

By grouping categories based on their associated analysis phases we see similar trends over time (Fig 3C). We see increases in the percentage of tools performing tasks in Phase 1 (quantification), across multiple phases (such as visualisation and dimensionality reduction) and alternative analysis tasks. In contrast the percentage of tools that perform gene identification tasks (Phase 4) has decreased and the percentage assigning cells (Phase 3) has remained steady. Phase 2 (quality control and filtering) has flucuated over time but currently sits at a similar level to when the database was first created. This too indicates a maturation of the analysis space as developers shift away from the tasks that were the focus of bulk RNA-seq analysis and continue to focus on those specific to scRNA-seq while working on methods for handling data from new protocols and performing alternative analysis tasks.

### Pipelines and toolboxes

While there are a considerable number of scRNA-seq tools that only perform a single analysis task, many perform at least two (Fig 3D). Some tools (dropEst [35], DrSeq2 [36], scPipe [37]) are pre-processing pipelines, taking raw sequencing reads and producing an expression matrix. Others, such as Scanpy [38], SCell [39], Seurat, Monocle and scater [40] can be thought of as analysis toolboxes, able to complete a range of complex analyses starting with a gene expression matrix. Most of the tools that complete many tasks are more recent (Fig 3E). Being able to complete multiple tasks using a single tool can simplify analysis as problems with converting between different data formats can be avoided, however it is important to remember that it is difficult for a tool with many functionalities to continue to represent the state of the art in all of them. Support for common data formats, such as the recently released SingleCellExperiment object in R [41], provides another way for developers to allow easy use of their tools and for users to build custom workflows from specialised tools.

### Alternative analyses

Some tools perform analyses that lie outside the common tasks performed on scRNA-seq data described above. Simulation is one alternative task that has already been mentioned but there is also a group of tools designed to detect biological signals in scRNA-seq data apart from changes in expression. For example alternative splicing (BRIE [42], Outrigger [43], SingleSplice [44]), single nucleotide variants (SSrGE [45]), copy number variants (inferCNV [46]) and allele-specific expression (SCALE [47]). Reconstruction of immune cell receptors is another area that has received considerable attention from tools such as BASIC [48], TraCeR [49] and TRAPeS [50]. While tools that complete these tasks are unlikely to ever dominate scRNA-seq analysis we expect to see an increase in methods for tackling specialised analyses as researchers continue to push the boundaries of what can be observed using scRNA-seq data.

## Discussion

Since October 2016 we have seen the number of software tools for analysing single-cell RNA-seq data almost triple, with more than 200 analysis tools now available. As new tools have become available we have curated and catalogued them in the scRNA-tools database where we record the analysis tasks that they can complete, along with additional information such as any associated publications. By analysing this database we have found that tool developers have focused much of their efforts on methods for handling new problems specific to scRNA-seq data, in particular clustering cells into groups or ordering them along a trajectory. We have also seen that the scRNA-seq community is generally open and willing to share their methods which are often described in preprints prior to peer-reviewed publication and released under permissive open-source licenses for other researchers to re-use.

The next few years promise to continue to produce significant new developments in scRNA-seq analysis. New tools will continue to be produced, becoming increasingly sophisticated and aiming to address more of the questions made possible by scRNA-seq data. We anticipate that some existing tools will continue to improve and expand their functionality while others will cease to be updated and maintained. Detailed benchmarking and comparisons will show how tools perform in different situations and those that perform well, continue to be developed and provide a good user experience will become preferred for standard analyses. As single-cell capture and sequencing technology continues to improve analysis tools will have to adapt to significantly larger datasets (in the millions of cells) which may require specialised data structures and algorithms. Methods for combining multiple scRNA-seq datasets as well as integration of scRNA-seq data with other single-cell data types, such as DNA-seq, ATAC-seq or methylation, with be another area of growth and projects such as the Human Cell Atlas [51] will provide comprehensive cell type references which will open up new avenues for analysis.

As the field expands the scRNA-tools database will continue to be updated with support from the community. We hope that it provides a resource for researchers to explore when approaching scRNA-seq analyses as well as providing a record of the analysis landscape and how it changes over time.

## Methods

### Database

When new tools come to our attention they are added to the scRNA-tools database. DOIs and publication dates are recorded for any associated publications. As preprints may be frequently updated they are marked as a preprint instead of recording a date. The platform used to build the tool, links to code repositories, associated licenses and a short description are also recorded. Each tool is categorised according to the analysis tasks it can perform, receiving a true or false for each category based on what is described in the accompanying paper or documentation. We also record the date that each entry was added to the database and the date that it was last updated.

### Website

To build the website we start with the table described above as a CSV file which is processed using an R script. The lists of packages available in the CRAN, Bioconductor, PyPI and Anaconda software repositories are downloaded and matched with tools in the database. For tools with associated publications the number of citations they have received is retrieved from the Crossref database (www.crossref.org) using the rcrossref package (v0.8.0) [52]. We also make use of the aRxiv package (v0.5.16) [53] to retrieve information about arXiv preprints. JSON files describing the complete table, tools and categories are outputted and used to populate the website.

The website consists of three main pages. The home page shows an interactive table with the ability to sort, filter and download the database. The second page shows an entry for each tool, giving the description, details of publications, details of the software code and license and the associated software categories. Badges are added to tools to provide clearly visible details of any associated software or GitHub repositories. The final page describes the categories, providing easy access to the tools associated with them. An additional page shows a up-to-date version of some of the analysis presented here with visulisations produced using ggplot2 (v2.2.1.9000) [54] and plotly (v4.7.1) [55].

### Analysis

The most recent version of the scRNA-tools database as of 2018-03-22 was used for the analysis presented in this paper. Data was manipulated in R (v3.4.3) using the dplyr package (v0.7.4) [56] and plots produced using the ggplot2 (v2.2.1.9000) and cowplot (v0.9.2) [57] packages.

## Declarations

### Ethics

Not applicable.

## Availability of data and materials

The scRNA-tools databases is publicly accessible via the website at www.scRNA-tools.org. Suggestions for additions, updates and improvements are warmly welcomed at the associated GitHub repository (https://github.com/Oshlack/scRNA-tools) or via the submission form on the website. The code and datasets used for the analysis in this paper are available from https://github.com/Oshlack/scRNAtools-paper.

## Competing interests

The authors declare no competing interests.

## Funding

Luke Zappia is supported by an Australian Government Research Training Program (RTP) Scholarship. Alicia Oshlack is supported through a National Health and Medical Research Council Career Development Fellowship APP1126157. MCRI is supported by the Victorian Government’s Operational Infrastructure Support Program.

## Authors’ contributions

LZ developed the scRNA-tools database and website. BP contributed to supervision of the project. AO oversaw all aspects of the project. All authors contributed to drafting the manuscript. All authors read and approved the final manuscript.

## Acknowledgements

We would like to acknowledge Sean Davis’ work in managing the awesome-single-cell page and producing a prototype of the script used to process the database. Daniel Wells had the idea for recording software licenses and provided licenses for the tools in the database at that time. Breon Schmidt designed a prototype of the scRNA-tools website and answered many questions about HTML and Javascript. Our thanks also to Matt Ritchie for his thoughts on early versions of the manuscript.

